# Differentiating the Physiological Signatures of Cochlear Synaptopathy and Inner Hair Cell Damage in a Chinchilla Model

**DOI:** 10.64898/2026.05.05.723072

**Authors:** Andrew Sivaprakasam, Ivy Schweinzger, Michael Heinz

## Abstract

Aging and noise over-exposure lead to complex mixtures of cochlear degradation that impair the structure and function of outer hair cells, inner hair cells (IHCs), and the cochlear nerve. However, IHC damage and cochlear synaptopathy (CS) remain pathologies “hidden” from the audiogram. This study aimed to identify and differentiate the physiological signatures of these two distinct pathologies using promising non-invasive assays: Envelope Following Responses (EFRs), Auditory Brainstem Response (ABRs), Wideband middle-ear reflexes (WB-MEMRs), and Distortion Product Otoacoustic Emissions (DPOAEs). We utilized chinchilla models of carboplatin-induced (CA) IHC damage (N = 4) and temporary threshold shift (TTS) noise-induced CS (N = 4) to compare the physiological signatures of each pathology. While both groups showed unchanged ABR thresholds two weeks after exposure, EFRs, ABR Wave V/I ratios, and MEMRs showed distinct effects of exposure. Despite non-elevated ABR-derived audiometric thresholds after exposure, both CA and TTS exposure resulted in severe in EFR “peakiness”, particularly for sharp, short-duty-cycle stimuli and significant elevations in ABR Wave V/I ratios. However, these findings were less-pronounced in the TTS-exposed animals. WB-MEMR amplitudes were decreased with elevated thresholds in both groups; this effect was more pronounced in the TTS group. Opposite trends in DPOAE amplitudes indicated that while both IHC damage and CS result in similar suprathreshold temporal coding deficits, effects on outer-hair-cell integrity and auditory efferent physiology may differ between the two pathologies. Future work and novel diagnostics should aim to distinguish these specific cochlear pathologies in clinical populations, or at the very least consider their overlap.

**Highlights:**

- A multi-metric diagnostic approach was used with chinchilla models of inner-hair-cell (IHC) damage and cochlear synaptopathy (CS).
- IHC damage and synaptopathy both cause suprathreshold deficits “hidden” from the audiogram.
- IHC damage results in more severe temporal envelope coding degradation than does synaptopathy.
- A combination of EFR “peakiness”, ABR Wave V/I ratio, and Wideband Middle Ear Muscle Reflex (WB-MEMR) appear to be useful measures for profiling IHC damage and CS.

## 1. Introduction

While remarkably robust, cochlear physiology is susceptible to a spectrum of degradation from natural phenomena like aging and noise exposure. Both presbycusis and hearing loss secondary to noise exposure are classically attributed to the loss of or damage to outer hair cells (OHC) in the Organ of Corti. However, relatively recent work in human temporal bones (Viana et al. 2015) and in mice (Sergeyenko et al. 2013) suggests that aging can also lead to cochlear synaptopathy (CS) - a decoupling of Type I auditory nerve fibers from their conjoining inner hair cell (IHC) synapses - which is accelerated by the addition of acoustic overexposure (Fernandez et al. 2015; Kujawa and Liberman 2009). CS has been highlighted as a significant, or even dominant, effect of moderate noise exposures that does not cause permanent elevations in auditory threshold—perhaps explaining the numerous cases of auditory perceptual deficits that remain idiopathic despite standard clinical workup (Tremblay et al. 2015). It is unlikely that CS always occurs purely in isolation, however. Indeed, several animal models indicate CS likely occurs with concomitant outer hair cell (OHC) damage after noise exposure (Kujawa and Liberman 2009; Valero et al. 2017; Pienkowski 2018). Therefore, isolating the effects of synaptopathy from OHC dysfunction, with the goal of a sensitive and specific diagnostic test, has recently become the focus of laboratory research studies (Vasilkov et al. 2021; Fernandez et al. 2020). However, the perceptual consequences of CS remain unclear. Leading hypotheses suggest that a primary sequela of CS is a disruption in the ability to understand speech in noise (Kujawa and Liberman 2015). This is partially supported by studies demonstrating correlations between speech in noise scores and measures thought to be indicative of CS; for example, ABR Wave I amplitude (Mepani et al. 2021; Megarbane and Fuente 2020) or subcortical envelope coding (Grant et al. 2020; Paul et al. 2017). Some studies report a lack of association between speech in noise scores and amplitudes of electrophysiological responses (Guest et al. 2018; Grose et al. 2017; Grinn et al. 2023; Lobarinas et al. 2017). Without a means of assessing cochlear micropathology *in vivo*, it remains quite unclear how specific and sensitive current assays of synaptopathy are in humans, and it is possible that these paradigms are not optimal (DiNino et al. 2022; Bramhall et al. 2019).

The impediment to measuring the consequences of synaptopathy in humans has been primarily due to using assays that do not contain specific measures of synapse function alone. Rather, these assays likely assume isolated synaptic damage, ignoring secondary damage (e.g., OHCs, IHCs, spiral ganglion cells) that is likely present, but difficult to assess without histology. This lack of specificity limits their clinical and research utility. Importantly, damage to the IHC body and stereocilia could present independent physiological deficits that manifest similarly (or differently) to the phenotypic presentation of isolated synaptic damage.

Animal studies using chinchillas have developed drug-exposure models that result in isolated IHC damage and loss. Carboplatin (CA) is effective in damaging IHC stereocilia and nuclei (Wake et al. 1994; Ding et al. 1999). Comprehensive peripheral physiological validations studies of this model by Axe & Heinz (2017) demonstrate that, while distortion product otoacoustic emissions (DPOAEs) and ABR thresholds remained intact post-exposure, suprathreshold Wave I ABR amplitudes were reduced. Histological data from this model demonstrated a 10-15% reduction in IHCs, with intact IHCs demonstrating substantial stereocilial damage (Axe 2017). Single-unit auditory-nerve responses from the same dataset demonstrated that, despite stable post-exposure ABR thresholds, the information available for the detection of modulations was reduced – contributing factors included 1) increased variability in temporal-coding metrics due to overall reductions in the total number of driven auditory-nerve-fiber spikes, 2) a reduction in the dynamic range of the IHC transduction function leading to impaired signal capture, and 3) a change in the apparent distribution of auditory-nerve-fiber spontaneous rates. Variability in IHC, OHC, and cochlear synaptic damage occurs post noise exposure or secondary to aging, which could cause a variety of perceptual and physiological deficits and potentially contribute to individual differences in human listener ability especially in noise. Therefore, a diagnostic assay or combination of assays that can differentiate these pathologies would have great clinical utility.

A number of other diagnostic assays show promise in this area. The Envelope Following Response (EFR) may capture subtle changes in electrophysiological processing that are not directly observed in ABR or DPOAE measurements. Chinchilla studies have demonstrated that permanent threshold shifts with OHC damage result in a low-frequency envelope over-representation in the EFR due to broadened tuning and distorted tonotopy confirmed by single-unit data (Parida and Heinz 2022, 2021; Henry et al. 2016; Zhong et al. 2014). This results in a paradoxical *increase* in EFR amplitude. Additionally, the EFR has been proposed as a helpful biomarker for detecting synaptopathy (Bharadwaj et al. 2019; Vasilkov et al. 2021; Encina-Llamas et al. 2019). These studies collectively imply that middle-aged listeners with normal hearing thresholds have reduced temporal coding abilities when compared to young, normal-hearing listeners, evidenced by *decreased* EFR amplitudes. This is presumably due to poorer phase-locking ability and decreased auditory-nerve synchrony across channels from synaptic loss or damage. The Wideband Middle-Ear Muscle Reflex (WB-MEMR) expands the clinically-utilized MEMR with a wideband elicitor, improving its potential sensitivity to synaptopathy in both chinchillas and humans (Bharadwaj et al. 2022). However, neither WB-MEMRs nor EFRs have been thoroughly explored in the context of IHC damage. One study demonstrated that clinically-standard MEMR amplitudes appear unaffected after IHC damage (Trevino et al. 2022), but no studies to the authors’ knowledge have similarly explored the EFR-related consequences of IHC damage and how they may compare to those found in CS.

The diagnostic utility of the EFR in identifying deficits like CS and IHC damage is stimulus dependent. Vasilkov et al. (2021) found that square-wave stimuli with short duty cycles better elicit subtle differences in EFRs from listeners with normal audiometric thresholds that could be at least partially attributed to CS, given correlations with age and noise-exposure status (Vasilkov et al. 2021). Specifically, a square-wave stimulus with 25% duty cycle and 95% modulation depth most reliably identified the three distinct populations tested (Young, Normal Hearing; Old, Normal Hearing; and Old, Hearing Impaired). In their study, ABR Wave V and I amplitudes did not separate these three groups quite as well, with significant overlap in amplitude values between the two older-listener groups. The authors suggest this was because ABR amplitudes elicited by 10 Hz click trains are more influenced by OHC-mediated cochlear gain prior to IHC transduction than 100 Hz rectangularly amplitude modulated (RAM) stimuli that fluctuate and have sharp envelope characteristics that are too temporally rapid for the cochlear gain function to impact. Theoretically, the isolation of cochlear synapse and IHC damage from OHC damage is then attainable—even more so when information from otoacoustic emissions can be incorporated (Keshishzadeh et al. 2021).

*But are the residual effects attributable to synaptopathy, IHC damage, or a mixture of both?* It remains an open, but critical, question whether synaptopathy alone is the driver of the effects detected with these diagnostics, or whether IHC damage plays a similarly important role. Better understanding the differences between these two unique pathologies would help guide our mechanistic hypotheses of why many listeners have “normal hearing” but struggle with auditory perception in complex environments. Both pathologies may be “hidden” from the audiogram and other common clinical diagnostics, but the EFR and WB-MEMR may present refined lenses through which to look.

The purpose of the present study was to utilize ABR, OAE, EFR, and WB-MEMR measurements to assess more precisely how the consequences of CS and IHC damage overlap and differ. It is currently unknown how comparatively specific or sensitive these assays are to either pathology, but it is generally hypothesized that they overlap in general effect and differ in severity (i.e. IHC damage presents as a “more severe” version of CS). The findings observed in these controlled chinchilla models of hearing loss help to identify potential additional physiological sources of the trends typically observed when using these assays in human subjects hypothesized to have “hidden” hearing loss. These precision diagnostic measures are commonly used to identify peripheral structures impacted by noise-exposure, ototoxic medications, aging, and genetics; thus, it is imperative to understand how they are affected by temporal coding deficits, particularly those caused by IHC damage.

## 2. Materials and Methods

Eight chinchillas (4 females, aged 32 to 48 weeks) were included in the study, separated into groups of 4 TTS (2 females), and 4 carboplatin (CA; 2 females) animals. No animals were used for previous experiments, and all procedures were approved by the Purdue Animal Care and Use Committee (#1111000123).

### 2.1 Exposures

#### 2.1.1 Carboplatin (CA) Exposure

Baseline data were recorded for four chinchillas (DPOAE, WB-MEMR, and EFR) prior to providing intraperitoneal CA injections (38 mg/kg) along with 10cc lactated Ringer’s solution as a supplement to offset any potential dietary changes during the initial adjustment period to the cytotoxic drug. Exposure procedures were identical to those described in the validation study by Axe & Heinz (Axe 2017). All CA animals were given soft foods and treats, were weighed, and given 10 cc of Ringer’s solution for four days following CA exposure to ensure that they maintained at least 70% of their pre-exposure body weight. CA animals were allowed to recover for two weeks prior to acquiring post-exposure data.

#### 2.2.2 TTS Noise Exposure

Four awake and unrestrained chinchillas were put into a metal cage housed in a wooden chamber (two animals were in the chamber during each exposure in separate cages) and were exposed to an octave-band noise (1 kHz center frequency) at 100 dB SPL for 2 hours. This method of exposing chinchillas to synaptopathic noise was identical to previous experiments (Bharadwaj et al. 2022), which showed both TTS and CS occurred following this noise exposure. Baseline measures (DPOAE, WB-MEMR, ABR, and EFR) were acquired prior to exposure. DPOAEs and WB-MEMRs were also recorded 1-day post-exposure to capture the temporary threshold shift, and two weeks post-exposure, after resolution of temporary noise-induced OHC dysfunction (Kujawa and Liberman 2009).

### 2.2 Awake Diagnostic Measurements (DPOAEs and MEMRs)

All chinchillas were put into a restraining tube, appropriate for body weight, which restricted their movement via a narrow opening for the neck, leaving the head exposed and their noses positioned in a customized nose-holder (methods adapted from Snyder & Salvi) (Snyder and Salvi 1994). A small mirror was placed beneath the nose allowed for monitoring of respiration during set-up. Animals were placed into a double-walled, sound shielded booth for testing. Respiration was further monitored using a video camera and pulse oximeter during awake recordings.

A microphone-transducer pair (ER10B+ probe, two ER-2 insert headphones; Etymotic, Elk Grove Village, IL, USA) with a foam tip was placed into the right ear of all animals for DPOAE and WB-MEMR recordings. DPOAEs were run using a 75/65 dB SPL (F1/F2) presentation level for frequencies from 500 to 10,000 Hz (F2 Frequency), with 6 logarithmically spaced steps per octave and a frequency ratio of 1.2. Four repetitions were conducted for each F2 frequency, each stimulus 2 seconds in duration with a 1-second inter-stimulus interval. DPOAE amplitudes were derived from the FFT magnitude of the average recorded ear-canal sound at the distortion product (2*F1-F2) frequency. The noise floor was estimated as the RMS of the signal contained within an octave below the distortion product frequency to an octave above the F2 frequency (excluding the DP, F1, and F2 frequencies). If the amplitude was less than 8 dB above the noise floor, it was discarded.

Following DPOAE recordings, the same microphone-transducer was used to record WB-MEMRs with click probe stimuli and broadband noise elicitor, consistent with studies investigating synaptopathic damage in mice (Valero et al. 2018), chinchillas (Bharadwaj et al. 2022), and humans (Mepani et al. 2020; Shehorn et al. 2020; Bharadwaj et al. 2022). Our WB-MEMR procedure was directly adapted from the validation studies described in Bharadwaj 2022 and Keefe et al. (Keefe et al. 2017). Briefly, one WB-MEMR block consisted of stimuli at 11 levels distributed evenly from 34 to 94 dB SPL with seven click probe and six broadband (0.5 to 8 kHz) elicitor presentations per level, adapted from Valero et al. (Valero et al. 2018). 32 blocks were run for each animal with an intertrial interval of 1.5 seconds. The elicitor was interleaved between 90 dB peSPL clicks and increased in 6 dB steps from 34 to 94 dB SPL. The reflex strength was quantified as the absolute change in absorbed ear-canal pressure spectrally averaged over a 0.5 to 2kHz region. This value increased with increasing elicitor level, and a growth function was estimated with a three-parameter sigmoid function for each animal. The growth function was baseline corrected using the mean reflex strength of the two lowest-level elicitors. The WB-MEMR threshold was quantified as the point where the function crossed 0.1 dB. One TTS animal was excluded from MEMR analyses due to substantial movement artifacts that limited an acceptable measurement SNR. Animals were allowed to recover for 24 hours following all awake recordings before any additional measures were collected.

### 2.3 Electrophysiological Recordings

#### 2.3.1 Anesthesia

Given the duration of electrophysiological testing and to reduce muscle artifact, animals were anesthetized. Sedation was induced with a subcutaneous dose of xylazine (4 mg/kg). A supplemental injection of ketamine (40 mg/kg) was administered subcutaneously 10 minutes later. 6 mL of lactated Ringer’s solution was delivered subcutaneously to keep animals hydrated throughout the experiment. Animals were then transported to a warm heating pad inside a double-walled, sound-attenuating booth where a homeothermic rectal probe-controller system was used to monitor body temperature. A pulse oximeter was placed on the animal’s paw, and the head was elevated above the torso using a bite bar (David Kopf Instruments). Eye ointment was applied to lubricate the eyes during sedation. Following the collection of all neurophysiological data (approx. 2 hours), atipamezole (0.2 mg/kg) was administered to reverse the sedation. The animal was carefully monitored until ambulatory and then returned to their housing and veterinary supervision. Animal recovery and weight were monitored for three days within the animal care facility.

#### 2.3.2 Calibration & Stimulus Presentation

To minimize inconsistencies in sound level due to natural variabilities in probe placement and ear canal resonance, an in-ear probe calibration was conducted during each experiment. Coefficients for a 256-tap digital inverse FIR filter were estimated from the frequency-response of the ear canal. The coefficients were then applied to a custom RX8 circuit (Tucker Davis Technologies) to ensure a flat (within +/- 2dB up to 10kHz) calibration in the ear. All stimuli were presented in the right ear through an ER-2 transducer coupled with the same probe used for otoacoustic emissions with the inverse filter applied.

Click and short duration (5 ms) 4 kHz tone burst stimuli were used to elicit ABRs. 1000 repetitions per polarity from 10 to 80 dB SPL in 10 dB steps were collected. An additional trial with stimuli presented within 5 dB of the visually identified threshold was conducted to improve precision. EFRs to stimuli with three different types of modulation were collected: square envelope with 25% duty cycle (SQ25), square envelope with 50% duty cycle (SQ50), and sinusoidal envelope (SAM). A carrier frequency of 4 kHz and modulation frequency of 100 Hz, with 100% modulation depth was used for all stimuli. All stimuli were 1.3 seconds in duration, with a .5 second interstimulus interval. 160 repetitions (80 per polarity) of each stimulus were collected at 80 dB SPL. To ensure a steady-state response free of onset-related effects, all EFRs were windowed from 0.6 to 1.3s prior to analysis.

#### 2.3.3 Recording Setup

For all electrophysiological recordings, a one-channel, vertical montage was employed in which the inverting electrode was placed behind the right pinna, tangential to the right mastoid. The non-inverting electrode was placed on top of the head, between the two tympanic bullae along the vertex of the interparietal bone. A grounding electrode was placed posterior to the nose, parallel to the nasal bone (Zhong et al. 2014). Physiological responses were pre-amplified using an ISO80 Isolated Differential Amplifier (World Precision Instruments) and then further filtered (0.3 to 3kHz) and amplified through a SR560 (Stanford Research Systems) amplifier for a total gain of 20k. All electrophysiological recordings were completed within the 2-hour window of sedation.

#### 2.3.4 Electrophysiological Analysis

ABR waveform processing scripts were developed in MATLAB. To estimate threshold, cross-correlations across sub-averages of all waveforms for a given level (1000 per/polarity, 2000 total) were computed. This approach is similar to the ABRpresto approach detailed by Shaheen et al. (Shaheen et al. 2024). Briefly, positive and negative polarity samples were divided into two groups of 1000 trials (500 trials/polarity each). Within each group, random sampling with replacement was used to select 800 waveforms (400 waveforms/polarity) for a subaverage. The resulting sub-averages for each group were cross-correlated. This procedure was repeated 200 times, to generate a distribution of cross-correlation values for each level and a sigmoid was fit to these data. Threshold was defined as the point on the sigmoidal curve 20% of the way to the inflection point. This method yielded similar (typically within 5 dB) threshold estimates as visual estimates by expert experimenters, with the added advantage of being more efficient and objective.

Wave I and V peak amplitudes and latencies were calculated for the loudest (80 dB SPL) click and 4k ABR responses using an automated dynamic time warping-based (DTW) approach. Four templates were generated from average responses, one for each exposure group at each time point (i.e. pre and post exposure). Wave I and V were identified on these templates, and the template was time-warped to the waveform from each animal. The resulting warping path was then used to identify Wave I and V on the individual waveform closest in Euclidean distance to the waves identified on the averaged template waveform. The identified wave was then constrained to the nearest local maximum to ensure a true peak. This method was more time-efficient than visual identification and particularly advantageous as it eliminated variability due to differences in experimenter judgment.

EFRs were spectrally analyzed by calculating the phase-locking value (PLV), a computation which quantifies the consistency with which summed neural units demonstrate a cohesive response phase across multiple repetitions of the eliciting stimulus. The PLV simplifies the analysis of the EFRs by allowing for a directly interpretable and unit-normalized value at a given frequency with an unbiased noisefloor. Trends in PLV and Fast-Fourier Transform (FFT) magnitude-based analyses are strongly correlated; however, PLV-based analyses are more sensitive to lower-amplitude, higher-frequency harmonics that contribute to the “peakiness” of responses important to this study (Zhu et al. 2013). Studies that use FFT magnitude often vary in how they compute noise floor and SNR criterion (Van Der Biest et al. 2023; Mepani et al. 2021; Vasilkov et al. 2021), which limits the generalizability and direct comparison of their findings to previous and future work.

Our implementation of the PLV estimate follows the calculations and guidelines provided in Zhu et al. (2013). In summary, for each individual animal, one fifth of the recorded trials from each polarity (16 trials/polarity, 32 total) were randomly drawn with replacement and averaged together. Phase as a function of frequency was computed using an FFT. This process was repeated 200 times to compute a distribution of phase as a function of frequency, and the phase locking value as a function of frequency was determined as the average unit vector strength across these repetitions (Fig 1). The resulting PLV spectrum contained several (typically ∼16) harmonics of the modulation frequency (100 Hz). To assess the relative strength of the PLV for high and low harmonics, we calculated R_PLV_, the sum of the upper harmonics relative to the lower two harmonics (Eq 1, Fig 1).

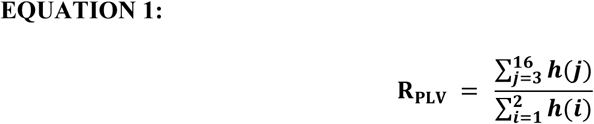

where *h* denotes the set of PLVs corresponding to each harmonic of 100 Hz in the EFR.

**Figure 1.**
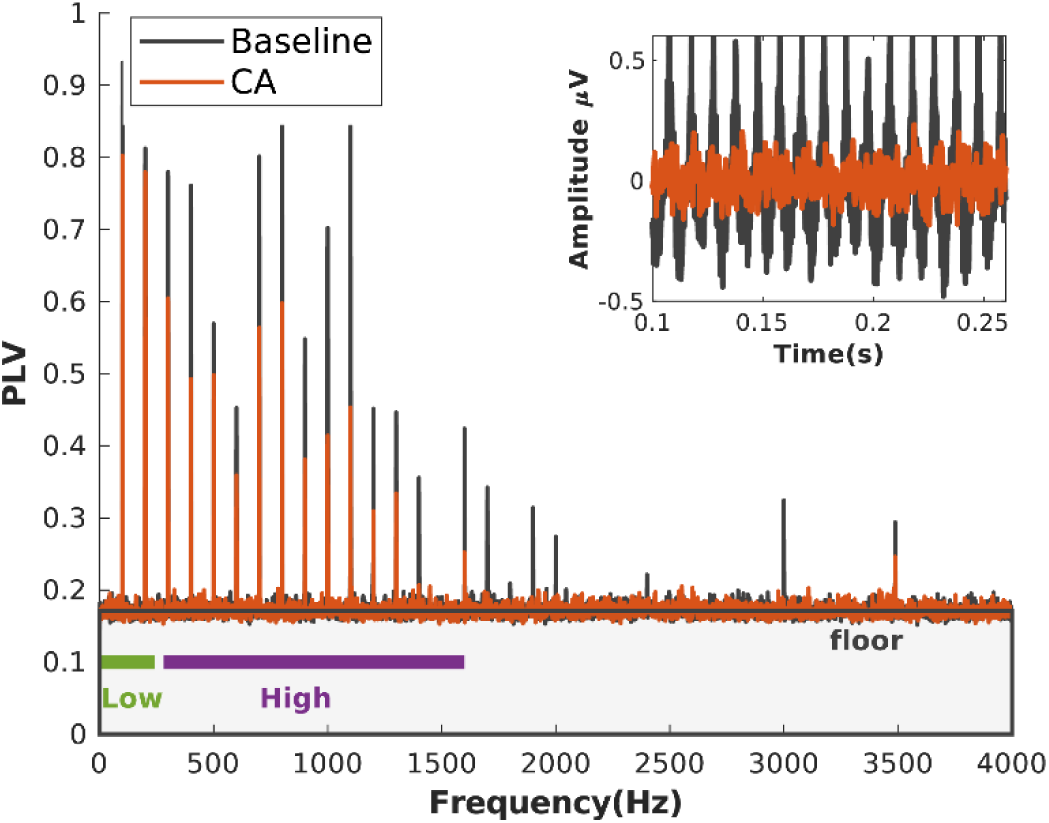
Averaged Phase-Locking Value (PLV) spectrum and time waveforms (upper right) of the EFRs recorded from carboplatin-exposed chinchillas pre (black) and post exposure (red). The signal energy carried within the upper harmonics (3 to 16) was normalized by that carried within the lower harmonics (1 and 2) to quantify changes in signal “peakiness”.

### 2.4 Statistical Analysis

All statistical analyses were performed in R (R Core Team, 2021). Linear mixed-effects models were used to infer the effects of exposure and group on the observed trends in the collected measures. Exposure group, time-point, and their interaction were modeled as fixed-effects terms, with individual animals modeled as a random effect. For TTS measures with 3 time points (baseline, 24 hours, and two weeks), the 24-hour time point was removed prior to statistical analysis when including CA data (baseline and two weeks). Separate statistical models were derived with baseline and 24-hour post-exposure time points for the TTS group. The models were derived using the lme4 library, with an F approximation for scaled type-II Wald statistic with Kenward-Roger degrees-of-freedom used to characterize the significance of the fixed effects (Bates et al. 2024). Post-hoc comparisons to investigate the significance of effects specific to a given group and exposure were conducted using estimated marginal means predicted from the linear mixed-effects models, using Tukey’s adjustment for multiple comparisons (Lenth et al. 2024). Simple linear models were used to identify correlations between unique measures (i.e. R_PLV_ vs ABR amplitude/latency).

## 3. Results

### 3.1 Auditory Brainstem Responses (ABRs)

Our selected TTS and CA exposure paradigms did not result in significant ABR threshold increases two-weeks after exposure, consistent with previous studies that used these exposure paradigms (Bharadwaj et al. 2022; Axe 2017). The 4-kHz thresholds for both groups showed no significant effect of group, exposure or their interaction. The CA click thresholds were not significantly different (at the single-group level) after exposure (Figure 2). There was, however, a modest but statistically significant decrease in click thresholds in the TTS group after exposure (mean difference = 10.6 dB SPL, t(6) = 3.769, p = 0.035, Tukey Adjusted), which contributed to a significant group-exposure interaction in the linear mixed-effects model (F(1,6) = 8.597, p = 0.026).

**Figure 2.**
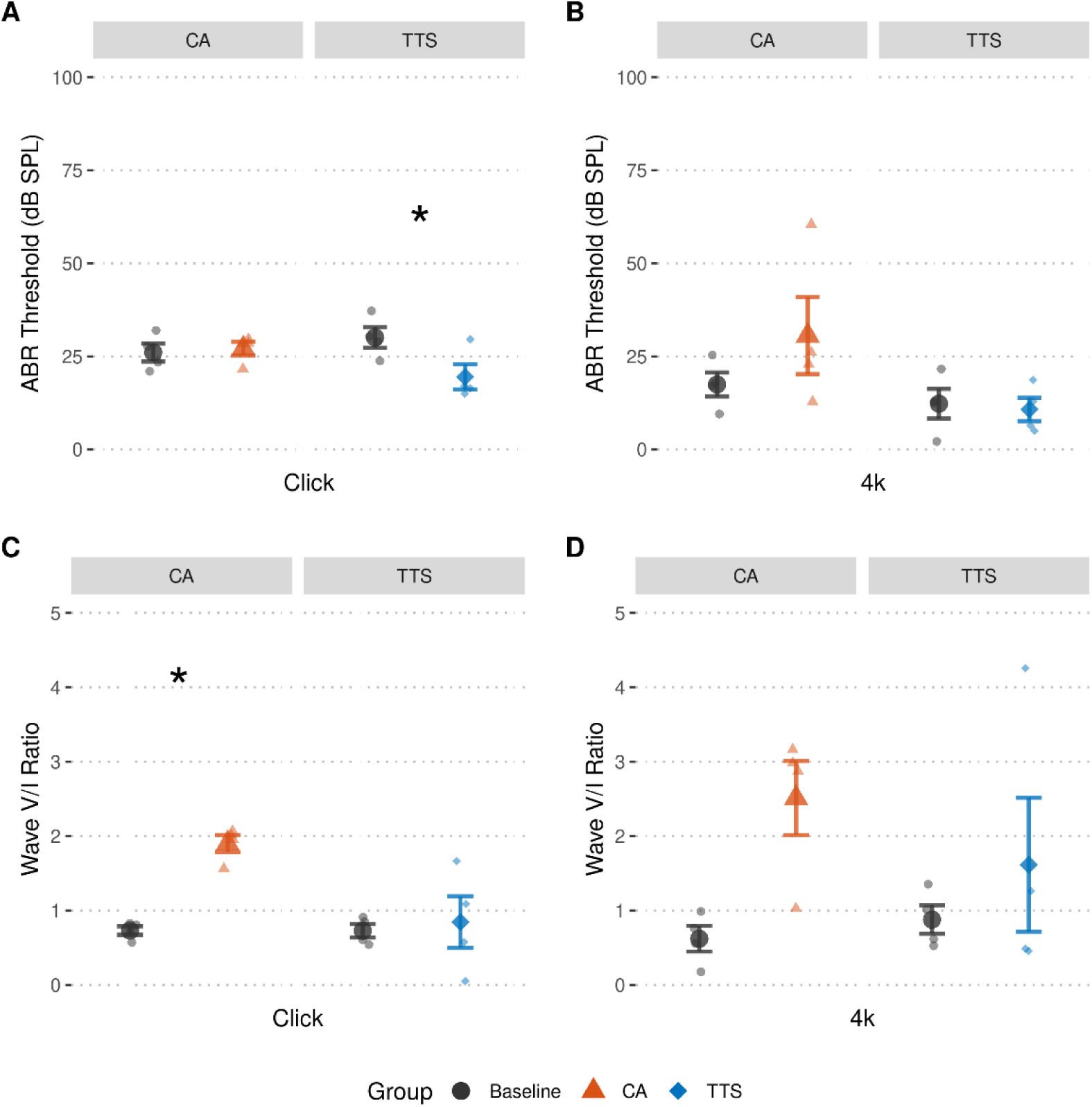
Temporary-Threshold-Shift (TTS) inducing noise and Carboplatin (CA) exposure do not impair ABR-derived auditory thresholds (A-B), but do result in elevated ABR Wave V/I ratios (C-D). Smaller points represent individual data, larger points represent the sample mean. Error bars represent SEM, * indicates p<0.05 at the single-group level. The main effect of Exposure across groups was significant (p<0.05) for changes in Wave V/I ratio.

Despite an absence of threshold elevations after exposure, changes in waveform morphology after both exposures were observed for both groups. These changes were most salient in the highest level (80 dB SPL) ABRs and when waveforms were averaged across animals (Figure 3), so only data at this maximal presentation level were used for these analyses. Grossly, Wave I amplitude reductions and Wave V amplitude increases were seen after either TTS or CA exposure, with CA exposure resulting in more defined effects (Figures 2 and 3). While changes in Wave I and Wave V amplitudes were not statistically significant at the single or multiple-group level, elevations in the Wave V/I ratio were more defined. The absolute values of Wave I and V amplitude were taken prior to computing this ratio to avoid occasional slightly negative values that occur when wave amplitudes are below the DC-corrected baseline. The Wave V/I ratio computed from click responses demonstrated a significant (F(1,6) = 11.462, p = 0.015) main effect of exposure, group (F(1,6) = 7.762, p = 0.032), and group-exposure interaction (F(1,6) = 7.678, p=0.032). The ratio increase was significant at the single-group level for CA-exposed animals (t(6) = -4.353, p=0.019, Tukey Adjusted). 4k waveforms showed a similar increase in Wave V/I ratio, with a significant main effect of exposure (F(1,6) = 7.015, p=0.0381). Again, this effect was stronger in CA-exposed animals, though not significant at the single-group level for either exposure group (Figure 2). Wave I and V latencies were generally longer for both groups after exposure, with the 4k Wave V latency showing a significant (F(1,6) = 8.178, p = 0.029) main effect of exposure, though this effect was not significant at the single-group level.

**Figure 3.**
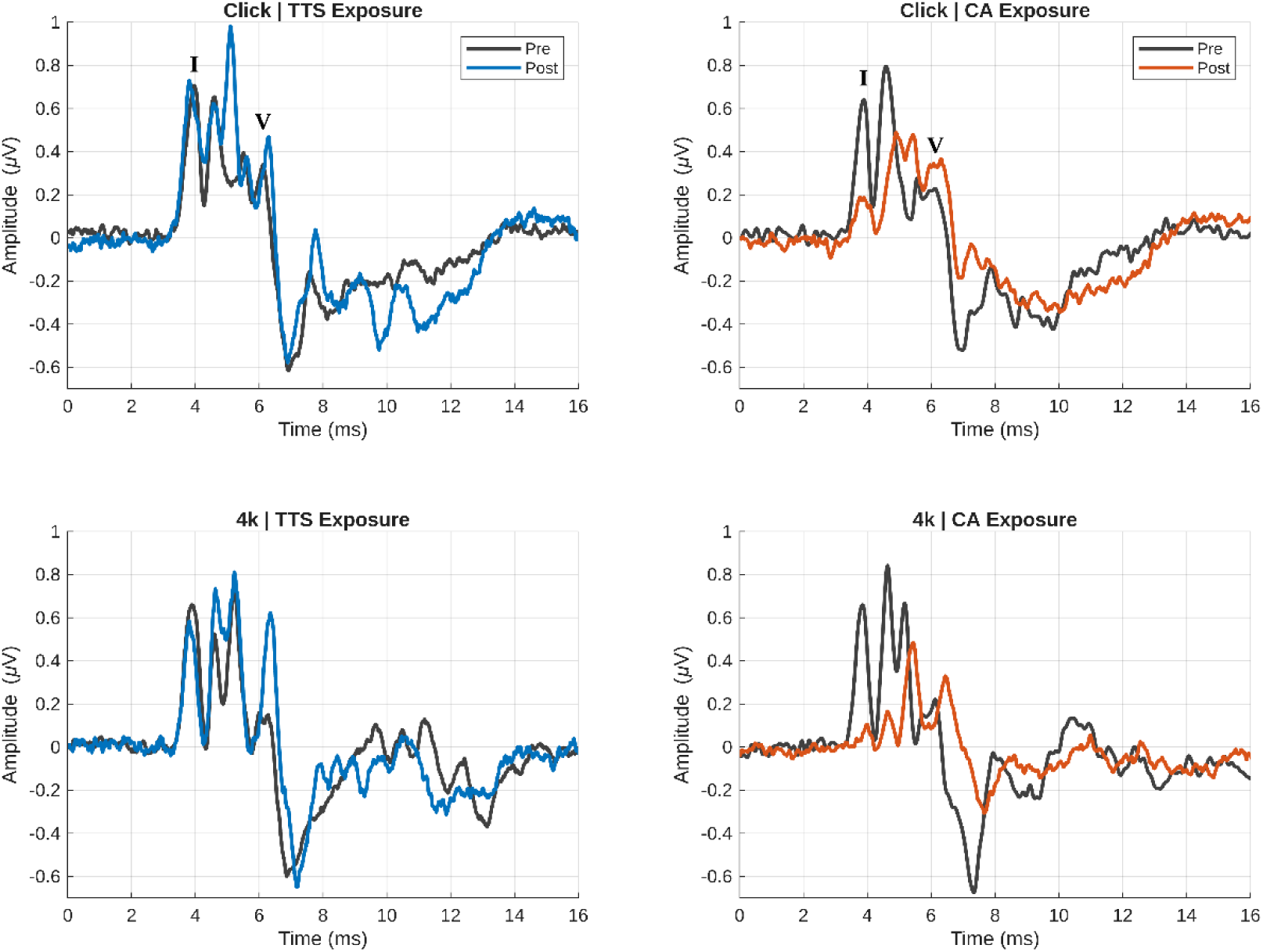
Morphological changes in grand-averaged ABR waveforms are evident after exposure to Carboplatin (CA) or a Temporary Threshold Shift (TTS) inducing noise. All waveforms are responses to a high-level (80 dB SPL) 4-kHz (Left Panel) or Click (Right Panel) stimulus. On average, Wave I amplitudes were reduced relative to Wave V. These findings were more salient in the CA group compared to the TTS group, with a longer latency after exposure, most notably observed in response to the 4-kHz tone.

### 3.2 Distortion Product Otoacoustic Emissions (DPOAEs) and Middle Ear Muscle Response Measures

DPOAEs were not reduced by carboplatin exposure, consistent with the expectation that this exposure primarily affects IHCs, sparing OHCs (Figure 4). However, we observed a mild elevation (2.79 dB SPL on average) of DPOAEs two weeks after CA exposure. This finding has been observed in a few other studies of IHC damage, potentially reflecting a reduced input to the efferent system and reduction of OHC-related gain control (Trevino et al. 2022; Axe 2017). In the TTS group, there was a transient significant (F(1,116.05) = 234.03, p<0.0001) reduction in DPOAE amplitudes (11.57 dB SPL lower on average) 24 hours post-exposure. DPOAE amplitudes mostly recovered (2.27 dB SPL lower than baseline on average) by the two-week post-exposure mark, similarly to ABR thresholds. Though the increase and decrease in DPOAE amplitudes after CA and TTS exposure, respectively, were modest in magnitude, they were consistent enough to lead to a significant (F(1,265.047) = 40.576, p<0.0001) Group*Exposure interaction.

**Figure 4.**
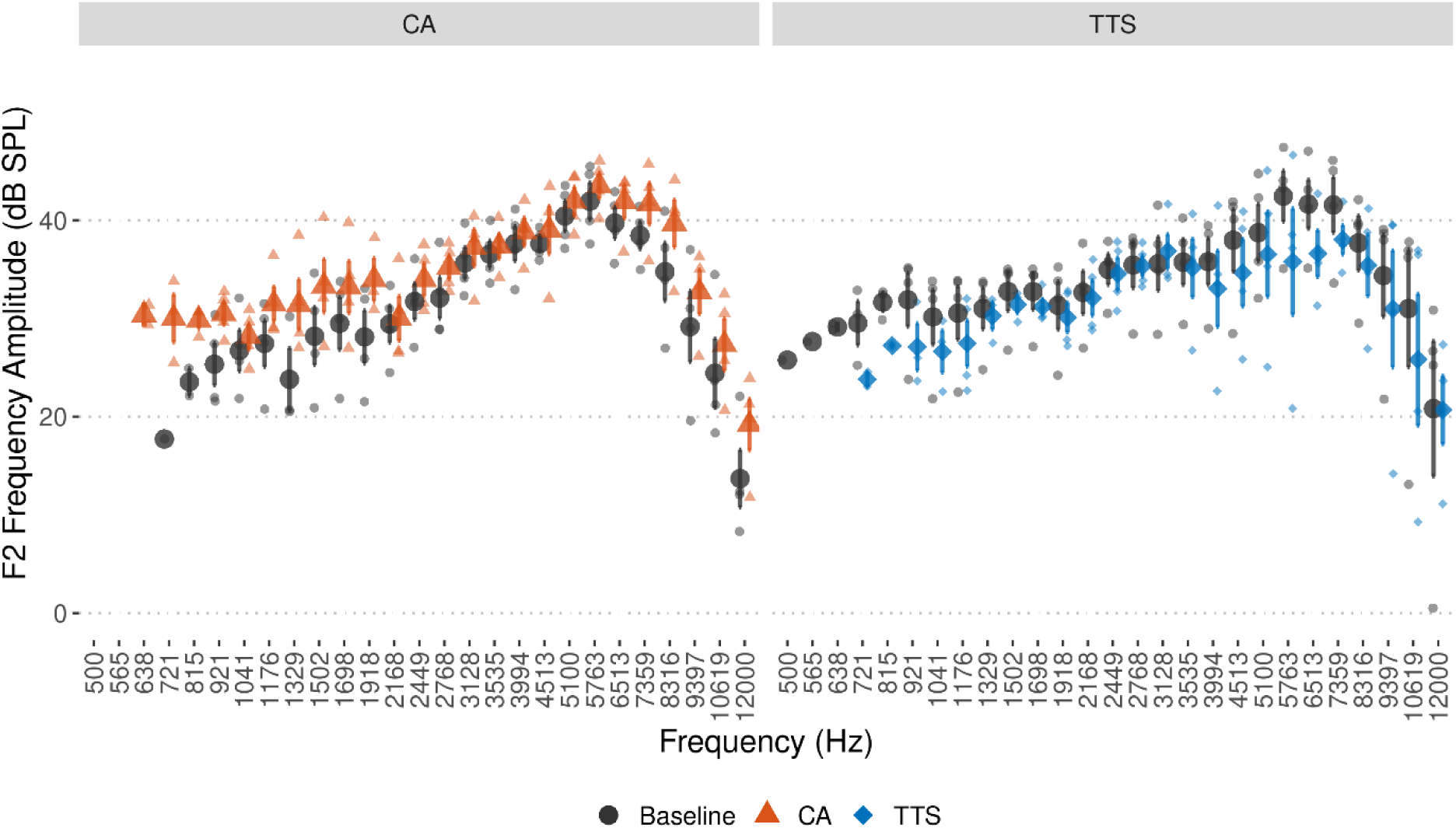
Distortion Product Otoacoustic Emissions (DPOAEs) remain stable after exposure to either Carboplatin (CA) or Temporary Threshold Shifts (TTS) inducing noise. Subtle DPOAE elevations and reductions are noted in the CA and TTS groups, respectively. These subtle and opposite findings were significant (p<0.0001), evidenced by a Group*Exposure interaction. Smaller points represent individual data; larger points represent the sample mean. Error bars represent SEM.

WB-MEMR thresholds were elevated in both groups two weeks after exposure (Figure 5B), with a significant main effect of Exposure (F(1,5) = 7.97, p=0.037), though these elevations were not significant at the single-group level. MEMR strength increased significantly with elicitation level, with a significant main effect of level (F(10,105) = 35.341, p<0.0001). After either exposure, there were significant reductions in MEMR strength (Figure 5A) at higher levels, though these reductions were more substantial in the TTS group: supported by a significant main effect of Exposure (F(1,105)=20.866, p<0.0001), and Group*Exposure (F(1,105)=11.4165, p=0.001), Level*Exposure (F(10,105) = 7.391, p<0.0001), and Group*Level*Exposure (F(10,105)=6.602, p<0.001) interaction effects. It should be noted that the pre-exposure WB-MEMR amplitudes (Figure 5A) were higher on average in the TTS group than the CA group, but this offset is mitigated by within-animal comparisons post-exposure. In the TTS group, thresholds were not significantly elevated 24 hours post-exposure (F(1,2) = 0.14, p=0.74); however, WB-MEMR strength was reduced at higher levels during this early timepoint (as it was two weeks post-exposure), supported by significant main effects of Level (F(10,42) = 11.980, p<0.0001), Exposure (F(1,42) = 6.560, p=0.014), and (Level*Exposure F(10,42) = 4.302, p=0.0003) interaction effects.

**Figure 5.**
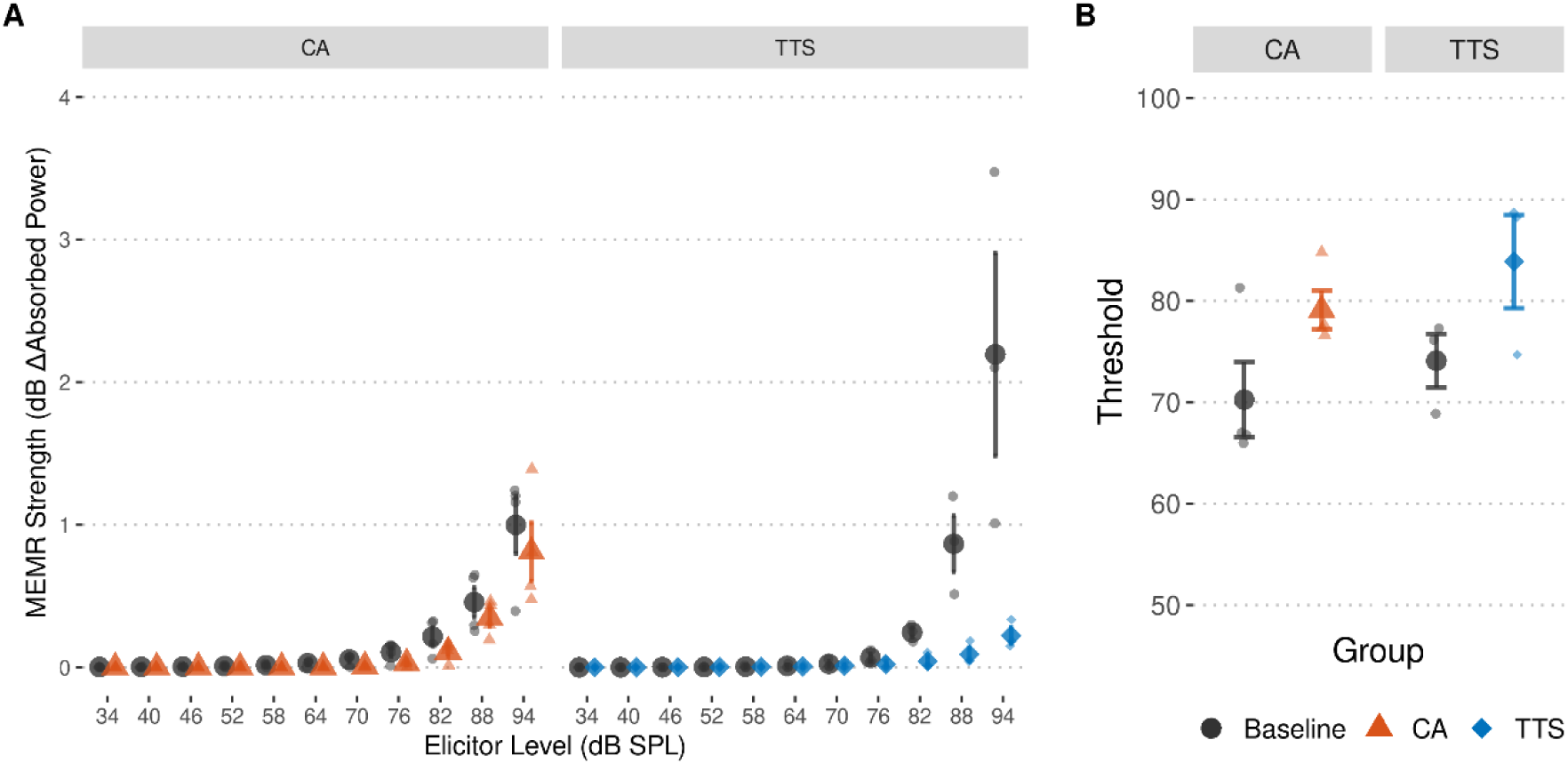
The Wideband Middle-Ear Muscle Reflex (WBMEMR) is degraded after either Carboplatin (CA) or Temporary Threshold Shift (TTS) inducing noise exposure. Reductions in MEMR strength are present, with more drastic effects in the TTS group. One TTS animal was excluded from these analyses due to significant movement artifacts. (A). MEMR Thresholds are similarly impacted across groups (B). Smaller points represent individual data; larger points represent the sample mean. Error bars represent SEM.

### 3.3. Envelope Following Responses

The temporal sharpness (or peakiness) of the EFR, particularly in response to square-modulated stimuli, was diminished in both CA and TTS-exposed chinchillas (Figures 6-7), but more notably in the CA group. The peakiness of the waveform presents spectrally as energy primarily in the upper (3rd through 16th) harmonics of the PLV spectrum. We quantified this effect as the sum of the PLV of upper harmonics relative to the lower two harmonics (Eq 1), R_PLV_. R_PLV_ was consistently reduced after carboplatin exposure in all three (SAM, SQ50, SQ25) stimulus conditions but was only noticeably reduced in response to the SQ25 stimulus after TTS exposure (Figure 8). In both cohorts, the square modulations (particularly SQ25) evoke a much stronger and peakier EFR at baseline compared to the sinusoidal modulation (SAM). The main effect of Exposure was significant (F(1,6)=8.412, p=0.027) for only the SQ25 stimulus condition, but approaching significance (F(1,6)=4.194, p=0.087) for the SQ50 stimulus, and not significant (F(1,6)=1.004, p=0.355) for SAM, demonstrating that the reductions in R_PLV_ were generally best observed in response to stimuli with sharper envelope and shorter duty-cycle. While these reductions were best-observed in the carboplatin group, they were not statistically significant in within-group comparisons of baseline to post-exposure measurements (i.e., contrasts of the estimated marginal means), likely due to a limited sample size. For the SQ25 R_PLV_ values, the main effect of the Group was significant (F(1,6)=8.4122, p=0.039), likely due to the slightly lower means in the carboplatin cohort before and after exposure compared to the TTS cohort.

**Figure 6.**
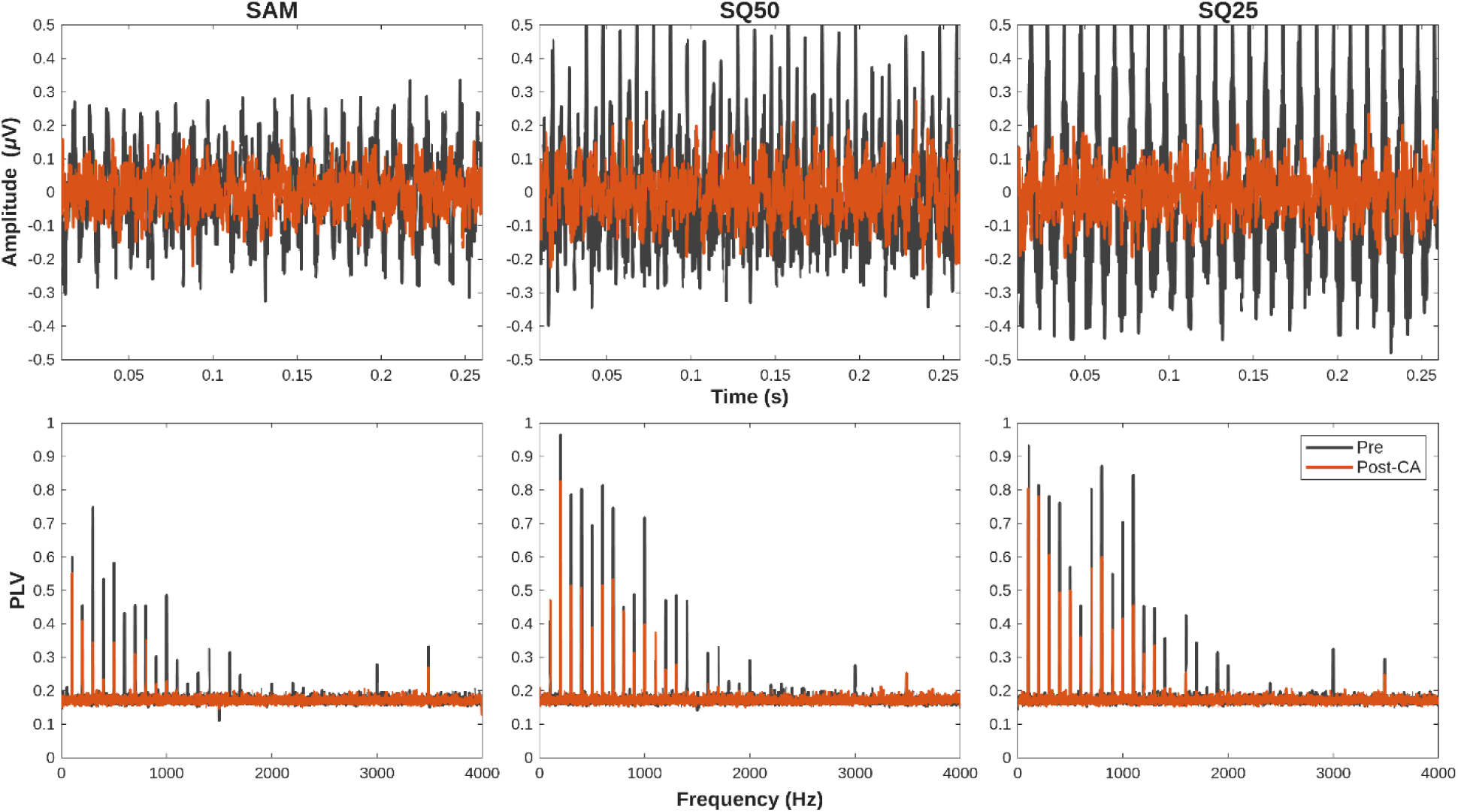
Grand-averaged EFR waveforms and PLV spectra from chinchillas before and after Carboplatin (CA) exposure. CA exposure reduces the temporal sharpness (“peakiness”) of EFRs to modulated stimuli (F_Mod_ = 100 Hz). Top: These changes are most noticeable in EFRs to stimuli with square modulation envelopes and 25% or 50% duty cycles (SQ25, SQ50). Bottom: Phase-Locking Values of the harmonics of the modulation were reduced after exposure, particularly those in the upper (3^rd^ to 16^th^) harmonics.

**Figure 7.**
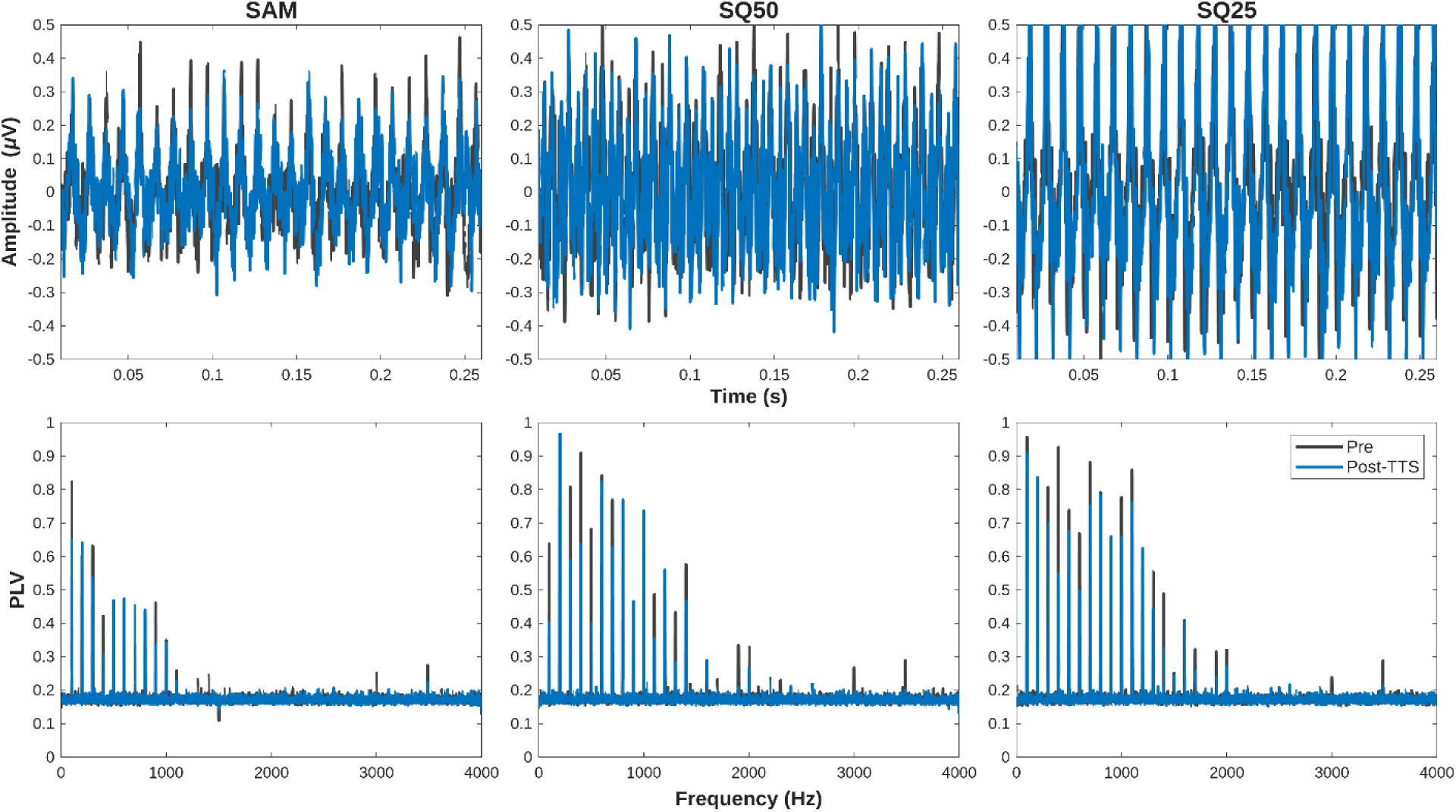
Grand-averaged EFR waveforms and PLV spectra from chinchillas before and after Temporary-Threshold-Shifts (TTS) inducing noise. TTS does not substantially impair EFRs to modulated stimuli (F_Mod_ = 100 Hz), evidenced by similarly strong temporal (Top) and spectral (Bottom) representations before and after exposure.

**Figure 8.**
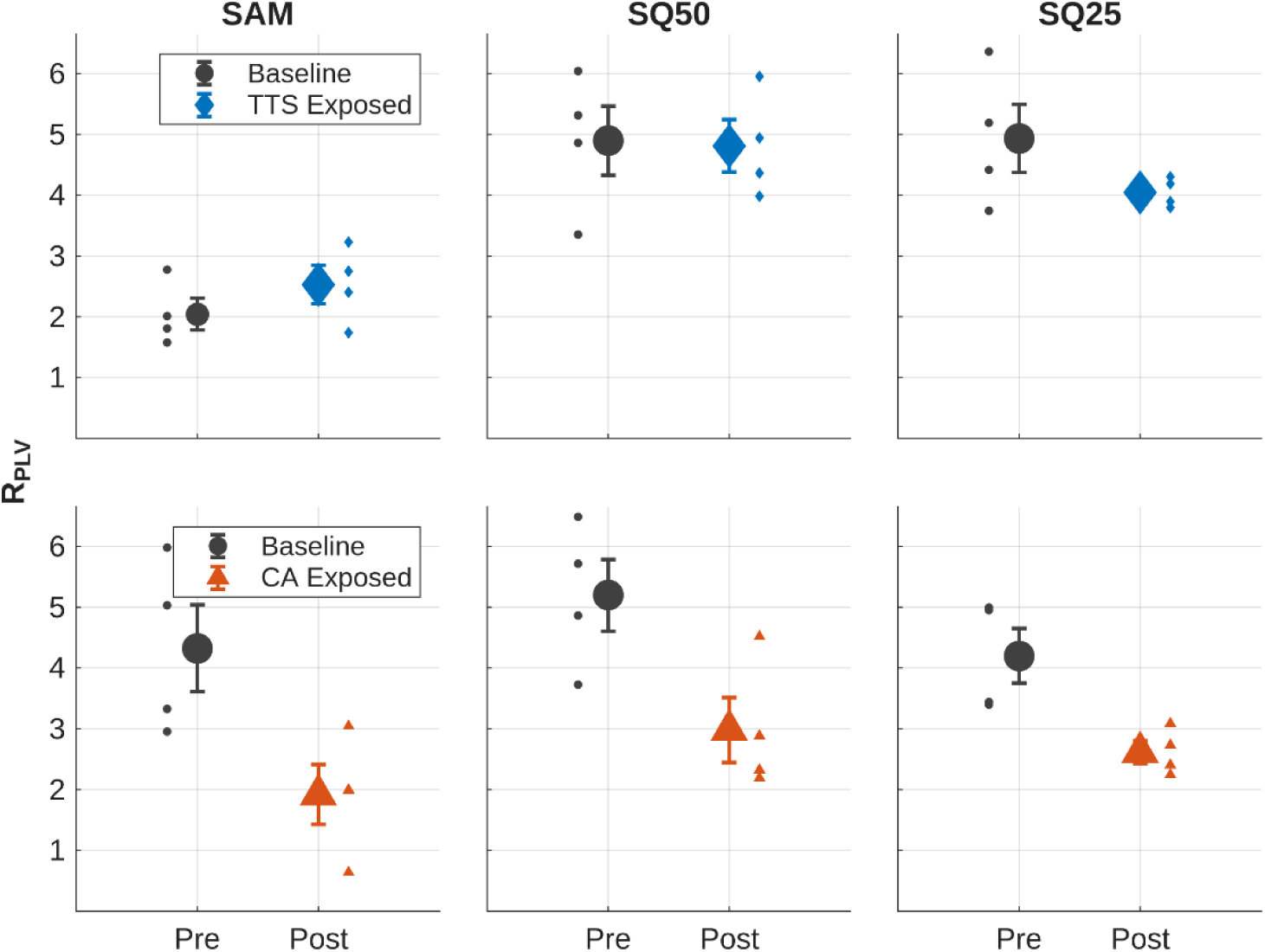
Changes in EFR waveform morphology after exposure were captured by the R_PLV_ metric, which captures the ratio of upper (3 to 16) harmonics to lower (1 to 2) harmonics. Animals in the TTS group (top) did not exhibit substantial changes in R_PLV_ after exposure (except for SQ25 stimuli). Animals exposed to CA (bottom) had notable reductions in R_PLV_ for all three stimuli. Thus, SAM and SQ50 stimuli showed different effects between CA and TTS exposures, while Sq25 showed reductions for both exposures. Smaller points represent individual data; larger points represent the sample mean. Error bars represent SEM.

Multiple regression analyses were also run to determine whether the diagnostic variables (e.g. ABR-based metrics, MEMR Strength, DPOAEs) were predictive of potential deficits in EFR PLV. Click and 4k Wave I ABR amplitudes were significantly correlated with R_PLV_ for SQ25 (Click | R = 0.63, p = 0.009; 4k | R = 0.76, p < 0.001; Figure 9) and SQ50 (Click | R = 0.54, p = 0.031; 4k | R = 0.54, p = 0.032) but not significantly correlated with R_PLV_ for SAM. Furthermore, R_PLV_ for SQ25, SQ50, and SAM were significantly negatively correlated with 4kHz-evoked ABR Wave V/I ratios (SQ25 | R = -0.56, p = 0.024; SQ50 | R = -0.66, p = 0.005; SAM | R = -0.55, p = 0.027), and both SQ25 and SQ50 were significantly negatively correlated with click-evoked ABR Wave V/I ratios (SQ25 | R = - 0.52, p = 0.037; SQ50 | R = -0.55, p = 0.027; SAM | R = -0.24, p = 0.37) (Figure 10). ABR Wave I and V latencies, DPOAE amplitudes, and MEMR thresholds were not significantly predictive of trends in these EFR stimuli. These findings are consistent with our expectations that upper harmonics in the EFR and ABR Wave I amplitudes are expected to be driven by auditory-nerve synchrony and should be reduced by either loss of intact synapses or IHC damage.

**Figure 9.**
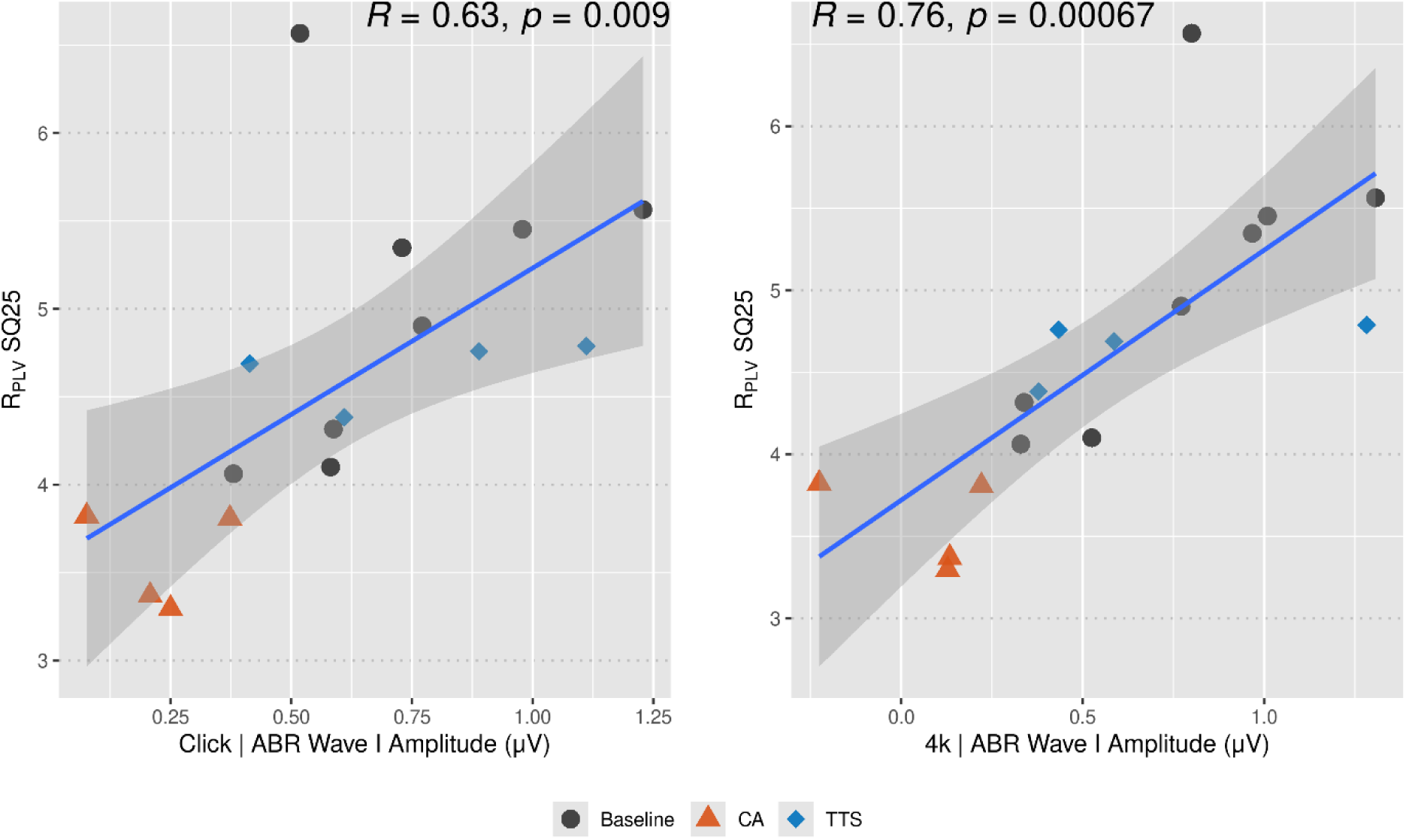
R_PLV_-quantified changes in EFRs to SQ25 stimuli were significantly correlated with ABR Wave I amplitudes for both high-level (80 dB SPL) click (right) and 4k (left) stimuli. Wave I amplitudes and R_PLV_ were most reduced after CA exposure, and remained relatively unchanged after TTS. Shading represents the 95% confidence interval of the linear regression estimate.

**Figure 10.**
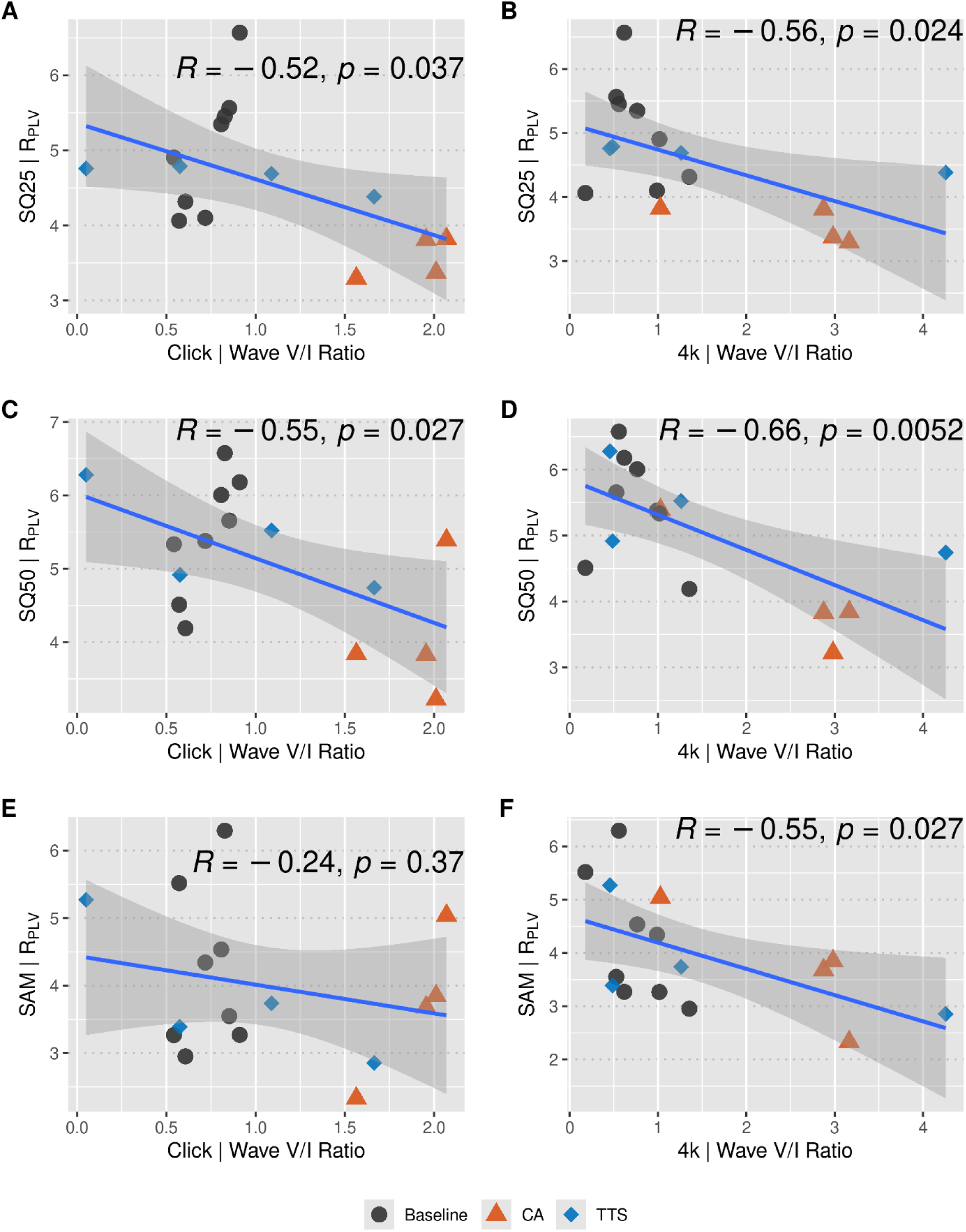
Click (Left) and 4k (Right) Wave V/I ratios were generally correlated with R_PLV_ for SAM, SQ50, and SQ25 stimuli. Reductions in R_PLV_ after TTS or CA exposure were significantly correlated with elevations in Wave V/I ratio, with the exception of the SAM vs Click pair (E).

## 4. Discussion

Outer-hair-cell integrity has been the primary focus of audiometric evaluations for decades; in part, due to the lack of agreed-upon, non-invasive, diagnostic assays of IHC and cochlear synapse function. However, the auditory system cannot function normally without an intact cochlear nerve and healthy IHCs. Hearing loss does not exclusively affect OHCs—histological temporal bone analyses from hearing-impaired human ears suggest the fraction of morphologically-intact IHCs is less than OHCs (Sayles and Heinz 2017). However, the physiological and perceptual consequences of IHC damage appear to be much more subtle. Indeed, while both IHC damage and CS could be considered “hidden” hearing loss pathologies given their limited effect on hearing thresholds, the latter has garnered more scientific attention and resources. Our present pilot work confirms that both types of hearing loss degrade the neural coding of stimulus modulations, even when audiometric thresholds are preserved.

An indicator of underlying pathology in this study was a reduction in temporal envelope coding ability, measured by EFR phase-locking values. We observed that high-frequency harmonics of the modulation frequency were typically reduced relative to the lower two harmonics, which we captured using the described ratio, R_PLV._ Contributions to higher harmonic (>300 Hz) frequency-following responses are expected to be dominated by peripheral structures like the auditory nerve and the cochlear microphonic from OHCs (Tichko and Skoe 2017). EFR metrics in this study focus on the polarity-tolerant component of this response (i.e., the *envelope*-following response) to reduce contributions from the polarity-sensitive cochlear microphonic. Therefore, R_PLV_ can be thought of as a ratio of auditory nerve to brainstem-dominant components of the EFR. Our findings then agree with the physiological expectation that IHC and cochlear-synapse dysfunction predominantly alter the neural representation of envelope by the auditory periphery. Any compensatory central gain would theoretically magnify differences in R_PLV_ after exposure.

The SNR of this measurement (and thus the power to resolve normal from impaired cochlear pathology) was highest when shorter-duty cycle (25% in this study) rectangular modulations were used, corroborating the developmental work on this stimulus by the Verhulst group (Vasilkov et al. 2021). Only in these sharply modulated stimuli were deficits in temporal envelope coding convincingly observed in TTS animals, our model of CS. However, animals with presumed IHC damage (exposed to CA) demonstrated more noticeable reductions in envelope coding fidelity across all stimuli. Given that ABR thresholds were not diminished in these groups post-exposure, *both* IHC damage and synaptopathy present with measurable temporal-coding deficits. The utility of rectangularly amplitude-modulated stimuli has been targeted towards identifying CS (Vasilkov et al. 2021; Van Der Biest et al. 2023; Ponsot et al. 2025; Verhulst and Vasilkov 2024), and previous works do not typically address the confounding or compounding effects of IHC damage. Our work demonstrates that these stimuli are partially sensitive, but not specific, to CS. In fact, we observed IHC damage appears to result in more drastic changes in temporal envelope coding. Future applications of these assays in research and in the clinic should acknowledge that the effects of IHC damage and CS on temporal envelope coding are difficult to separate, and that a multi-metric approach or combination of cochlear-focused diagnostics may be necessary to confidently attribute damage to either structure with specificity.

Indeed, the more common diagnostics of SNHL used in this study (ABR, DPOAE, MEMR) also appeared sensitive to the effects of suspected IHC damage and TTS. ABR Wave V/I ratios were significantly increased after exposure, indicating a potential disruption in temporal representation by the auditory periphery in both TTS and CA-exposed animals. These Wave V/I ratio increases were correlated with reductions in R_PLV_, particularly when sharply modulated EFR stimuli (SQ25, SQ50) were selected—further supporting this idea that both CS and IHC dysfunction result in temporal coding deficits in the auditory periphery and possibly compensatory central gain (Monaghan et al. 2020; Schrode et al. 2018; Salvi et al. 2017; Chambers et al. 2016). These waveform morphological changes were more severely altered in CA-exposed animals compared to TTS animals, though these differences did not always reach statistical significance (likely due to our small sample size). Longer wave latencies (particularly Wave V) and less qualitatively “peaky” waveforms were typically observed.

Interestingly, CA and TTS exposure resulted in marginal, but opposite, trends in DPOAEs. While TTS exposure resulted in a slight decrease in DPOAE amplitude, particularly at high frequencies, CA exposure resulted in subtle *increases* in DPOAE amplitude, leading to significantly different effects of exposure type. This paradoxical elevation in DPOAE amplitude after CA-induced IHC damage is not unique to our study. While the effect proved to be a differentiating factor in our statistical analysis, other studies have noted this increase to be of mixed presence and significance (Trevino et al. 2022; Axe 2017; Lobarinas et al. 2017; Trautwein et al. 1996; Wake et al. 1994). A possible mechanistic explanation for this is an IHC damage-induced reduction in cochlear afferent input to the central auditory system and subsequent disinhibition of OHC-dependent cochlear gain due to disruptions in auditory efferent function. Conversely, the TTS protocol could result in subtle OHC dysfunction that persists even after the temporary threshold shift and is not detectable by ABR-based assays.

Wide-band middle-ear muscle reflex thresholds were elevated, and the reflex strengths were reduced after either exposure. Reduced afferent input due to either IHC damage or CS could result in an impaired MEMR, despite normal thresholds. Indeed, the sensitivity of this metric is what makes it a promising tool for identifying these suprathreshold deficits (Bharadwaj et al. 2022; Hickox et al. 2017; Bharadwaj et al. 2019). The MEMR-related consequences of CS and IHC damage should theoretically be similar; however, the MEMR was markedly more reduced in the TTS group compared to the CA group. Indeed, similar preservation of the MEMR after IHC damage was observed by Trevino et al. (Trevino et al. 2022). One possible explanation is that synapse damage after TTS is thought to affect primarily high threshold, low-spontaneous-rate fibers (Furman et al. 2013; Bharadwaj et al. 2014). If changes to the MEMR are specifically driven by degradations to these fibers, it is possible that IHC damage alone would not cause a substantial disruption to low-spontaneous-rate fibers’ afferent signal. An alternative explanation is that TTS may also cause modest OHC dysfunction (e.g. uncoupling of stereocilia from the tectorial membrane) in the region responding to the elicited noise exposure (Nordmann et al. 2000). A potential consequence of this often-overlooked structural damage could be reduced input to the afferent limb of the MEMR pathway. Given the slight reductions in DPOAE amplitude we observed even two weeks after exposure, this remains a strong possibility. However, more work is needed to fully characterize why these two models of hearing loss differ in their MEMR consequences and whether these can be linked to changes in cochlear anatomy—given the limited sample size and absence of histological data in this initial study.

Collectively, the data presented here confirm that IHC damage and CS both result in suprathreshold deficits. The consequences of these types of damage are in contrast with those of OHC damage, which are studied more extensively in the current literature. IHC damage and CS appear to substantially affect temporal coding deficits, demonstrated primarily as changes in ABR and EFR responses. Notably, a degradation in temporal precision resulted in less “peaky” envelope representation, particularly in animals with presumed IHC damage. Conversely, OHC damage results in a lack of spectral precision and, in severe cases, distorted tonotopy (Henry et al. 2016; Parida and Heinz 2022). This reduced spectral precision results in phase-locking to envelope across a broader range of the cochlea, and an *enhanced* temporal-envelope representation (Zhong et al. 2014). Furthermore, OHC damage typically presents with more obvious threshold deficits and reductions in DPOAEs, which has facilitated its diagnosis and study. However, histological animal (Sayles and Heinz 2017) and human (Wu and Liberman 2022) temporal bone analyses suggest that IHC stereocilia damage could be similarly, if not more, prevalent in noise-exposed and aging population. Therefore, further exploration of the physiological and perceptual consequences of IHC damage is warranted.

No single measure utilized in this study could reasonably tease the consequences of IHC damage and CS apart from each other with diagnostic specificity. In most cases, CA and TTS animals demonstrated similar deficits that differed in their severity. Therefore, the combination of several measures may be more useful in separating these two pathologies. The modulation-type specificity of R_PLV_ for TTS but not CA exposures provides a potential avenue for future study to differentially diagnose these two subtypes of SNHL and may result from stereocilial effects on the on the nonlinear IHC transduction function following CA, but not TTS exposures. Also, changes in DPOAEs, though subtle, were opposite in CA and TTS animals. Pairing these changes with ABR, EFR, and WBMEMR measures could help differentiate these two pathologies. A characterization of the specificity and sensitivity of each of these measures, and their joint utility, could be more effectively done with a larger dataset that includes a wider array of exposures and cochlear histology. The present work presents early evidence that IHC damage is a significant form of “hidden” hearing loss with consequences overlapping, but in some important ways distinct from, those of CS. Future work must consider IHC damage in addition to CS as a possible source of suprathreshold deficits.

## Acknowledgements

A.S. and I.S. designed and performed the experiments, analyzed the data, and provided an initial draft of the paper.

M.H. assisted with study design and provided critical feedback on experimental techniques and the manuscript. The authors also wish to thank Dr. Samantha Hauser for her feedback on an early version of the manuscript. Research funding was supported by NIH NIDCD F30DC020916 (A.S.), 1T32DC016853 (A.S.), R01DC009838 (M.H.).

## Data Statement

All raw data collected may be found on Zenodo (DOI 10.5281/zenodo.20025719). Pre-processing, processing, and statistical analysis code is available on GitHub (https://github.com/sivaprakasaman/Sivaprakasam_Schweinzger_CA_TTS_Code).

## Notes

https://github.com/sivaprakasaman/Sivaprakasam_Schweinzger_CA_TTS_Code

https://zenodo.org/records/20025720

### Competing Interest Statement

The authors have declared no competing interest.

